# Mucin-laden PLGA nanoparticles inhibit neutrophil extracellular trap formation

**DOI:** 10.64898/2026.11.20.766221

**Authors:** Allison Boboltz, Owen Tabah, Alexis Johnson, Gregg Duncan

## Abstract

Neutrophils mount antimicrobial and pro-inflammatory responses to infection and tissue injury through degranulation, phagocytosis, and release of neutrophil extracellular traps (NETs) generated via a process also known as NETosis. However, excessive or prolonged neutrophil activation can damage host tissues and accelerate disease progression. As such, NETs have emerged as an important target in therapeutic development where nanoparticle-based strategies have been pursued. However, nanoparticles made of poly(lactic-co-glycolic acid) (PLGA), one of the most widely used biodegradable polymers for drug delivery, have been shown to promote NETosis which may compromise its use in neutrophil-driven disease states. Considering this, we sought to leverage the innate immunomodulatory properties of mucins to mitigate PLGA nanoparticle-induced NETosis. Specifically, we developed a new formulation of mucin-functionalized PLGA nanoparticles using commercially available porcine gastric mucin (PGM). We found PGM-laden PLGA NP significantly reduced the frequency of NETosis *in vitro* in activated neutrophil-like HL-60 cells and primary human neutrophils, down to levels observed in unstimulated cells. Importantly, this inhibitory function was observed when PGM-laden PLGA NP were administered to neutrophils either prophylactically or therapeutically up to 1 hour after NET activation. We also observed PGM-laden PLGA NP were readily internalized in neutrophil-like HL-60 cells to enable intracellular delivery of therapeutic cargoes. Taken together, this bioinspired approach offers a promising strategy to improve PLGA-based nanomedicines for the treatment of neutrophilic inflammatory diseases with the dual functionality of mitigating excessive NETosis while simultaneously facilitating the delivery of therapeutic cargoes.

## Introduction

Neutrophils are the most abundant circulating leukocyte in humans, comprising approximately 50-70% of the total immune cell population^1^. Neutrophils employ multiple killing mechanisms including phagocytosis, de-granulation, and the release of neutrophil extracellular traps through a process known as NETosis^1^. Neutrophil extracellular traps (NETs) are web-like complexes of de-condensed chromatin and entangled granular proteins which can effectively entrap and degrade pathogens. However, excessive NETosis can cause further downstream inflammation and directly harm native tissues due to increased exposure to anti-microbial granular proteins and histones. Numerous inflammatory conditions including autoimmune diseases, cancers, and infections involve excess NETosis that contributes significantly to disease pathogenesis.

Poly(lactic-co-glycolic acid) (PLGA) is one of the most widely used and studied synthetic polymers for drug delivery applications, included in clinically approved treatments for inflammatory conditions. Its widespread use in nanoparticle formulations can be attributed to its desirable properties such as biodegradability and high encapsulation efficiencies of various drug types^4^. As of 2025, PLGA is a major component in over 25 long-acting injectable drug products approved by the United States’ Food and Drug Administration (FDA). PLGA-based particles are the most common formulation type, with ∼15 FDA-approved products^6^. However, prior research demonstrated that unmodified PLGA nanoparticles (NPs) induce extensive NETosis *in vitro* using primary human polymorphonuclear leukocytes (PMNs)^7^. This propensity of PLGA to trigger NETosis may induce unwanted reactions that undermine their therapeutic use, particularly in diseases with neutrophilic inflammation. Many inflammatory conditions associated with increased NETosis overlap with those for which PLGA-based drug delivery formulations are being developed, including endometriosis^8^, periodontitis^9^, type 2 diabetes/diabetic complications^10^, and cancer^11^.

Using a bio-inspired approach from the human mucosa, we hypothesized PLGA NPs could be engineered that are intrinsically suppressive of NETosis. Specifically, recent work from our group has shown that airway mucin glycoproteins can strongly inhibit NETosis^12^. Mucins are large, heavily glycosylated proteins that are both secreted to form mucus and are expressed as cell-membrane bound proteins^13^. Mucins interact with immune cells via their associated glycans and serve a variety of immunomodulatory functions^14,15^. In addition to airway mucins, bovine cervical mucins have been shown to inhibit NETosis via sialic acid glycans^16^. Given the anti-NET activity of mucins via sialic acid, we sought to translate these prior findings to address the potential limitations of PLGA particle formulations used in disease indications involving neutrophilic inflammation.

The research presented herein provides evidence supporting the use of a bioinspired drug delivery approach to limit NETosis in inflammatory diseases. Prior work has investigated alternative synthetic polymers such as salicylic acid-based nanoparticles for the inhibition of NETosis^7^. Sialic acid-conjugated latex beads have also shown efficacy in preventing NETosis *in vitro*^17^. In addition, porcine gastric mucin-based nanoparticles have been developed and are biocompatible both *in vitro* and *in vivo*, but have not been tested for NETosis applications^18,19^. Herein, we sought to utilize porcine gastric mucins to modify the immunomodulatory function of conventional PLGA nanoparticles which may limit their use as agents in neutrophil-mediated inflammatory diseases. In doing so, we found that adding mucins to PLGA nanoparticle formulations ameliorated NETosis without sacrificing their drug delivery potential, which could enhance the efficacy of therapeutics for diseases in which neutrophilic inflammation is prevalent.

## Methods

### Nanoparticle Synthesis

Before particle preparation, porcine gastric mucins (PGM) (Sigma Aldrich; Type III, bound sialic acid 0.5-1.5%) were dissolved in ultrapure water with stirring at room temp for 20 minutes and PLGA (Polysciences, Poly(D,L-lactide-co-glycolide) 50:50 acid-terminated) was dissolved in chloroform. To make particles with varying concentrations of PGM, 1ml of 0, 1, 2.5, or 5 mg/ml PGM and 100 µL of 15 mg/ml PLGA was added to a microcentrifuge tube. The tube was placed on ice and probe sonicated (Qsonica Q125) into a homogenous emulsion at 50% intensity for 3 minutes on a 1 second on and 4 seconds off cycle. After sonicating, the tubes were placed on a rack and shaken at 180 rpm for 1-2 hours at room temp to evaporate chloroform. The nanoparticle solution was centrifuged at 1000g for 3 min to pellet any debris. 800 µL of supernatant was removed and filtered through a 0.45 µm syringe filter into a clean tube to remove contaminates.

### Physicochemical Characterization of Nanoparticle Size, Surface Charge, and Stability

Nanoparticle solutions were diluted 1:1000 in 10 mM NaCl and particle hydrodynamic radii (reported as the Z*-*average) and zeta potential was determined by dynamic light scattering (DLS) with a Malvern Zetasizer. To test nanoparticle stability in an ionic solution, 10x PBS was diluted in the particle solutions then automated measurements were performed every 15 minutes for 12 hours. Additionally, nanoparticle solutions were diluted 1:1000 in ultrapure water and analyzed using a ZetaView nanoparticle tracking analysis (NTA) system to evaluate particle concentration.

### DiD Loading and Efficiency

For subsequent cell uptake analysis, nanoparticles were loaded with DiD (Thermo Fisher D7757) as a fluorescent probe. DiD was added to the 15 mg/ml PLGA solution at a 0.4 wt:wt ratio with the PLGA. Then, nanoparticle synthesis was performed as mentioned above. To characterize the loading in the final solutions, PLGA and PGM-PLGA nanoparticles loaded with DiD were diluted 1:10 with DMSO to dissolve the PLGA and release the DiD. Fluorescence intensity of each final solution was measured with a plate reader (excitation/emission: 600/668). The concentration of the extracted DiD was determined by comparison to a standard curve of DiD dissolved in DMSO at a maximum concentration of 10 µg/ml with five serial dilutions down to 0 µg/ml.

### Flow Cytometry

500 µL of dHL-60 cells at a density of 10^6^ cells/mL was added to the wells of a sterile 24 well plate. Cells were incubated with 100 µL of either DiD loaded PLGA or DiD loaded PLGA-PGM (stock concentration of 5 mg/mL) NPs for 30 minutes prior to adding 100 nM PMA or a vehicle to each well. Cells were incubated for 1 hour following treatment with PMA or vehicle (PBS). The cells were stained with Sytox green diluted to 0.05 µM for 15 minutes to assess cell viability. The cells were then collected in microcentrifuge tubes and washed twice with PBS. The samples were analyzed with a BD FACSCelesta flow cytometer. We gated to exclude dead cells (Sytox green positive) and assess the DiD fluorescence (indicating NP uptake) of live cells.

### Confocal Microscopy

250 µL of dHL-60 cells were seeded in chamber well slides at a concentration of 10^6^ cells/mL. 50 µL of DiD-loaded PLGA NPs, PGM PLGA NPs (5 mg/mL PGM stock), or blank PBS (no NP treatment control) was added to each well and incubated for 30 minutes at 37°C and 5% CO_2_. PMA (100 nM final concentration) or a vehicle was added to each well, then dHL-60 cells were incubated for 4 hours. Cells were stained with Sytox green diluted to a final concentration of 0.05 µM for 15 minutes and washed 3 times with PBS. Cells were fixed with 4% PFA, then each slide was sealed with a coverslip. Slides were imaged at 10x magnification using a Zeiss LSM confocal microscope.

### HL-60 Culture and Differentiation into dHL-60 Cells

The immortalized human acute myeloid leukemia cell line HL-60 (ATCC CCL-240) was grown in RPMI 1640 media supplemented with 20% heat inactivated fetal bovine serum and 1% penicillin-streptomycin (complete RPMI 1640 media).^28^ The HL-60 cells were cultured for at least 1 week prior to inducing differentiation to achieve a neutrophil-like state. Media was replaced every 2-3 days and cells were maintained at a density between 10^5^ and 10^6^ cells/mL. To differentiate HL-60 cells, 1.3% DMSO was added to the complete RPMI 1640 media for a period of 5-7 days.^28,53^ The DMSO-containing complete RPMI 1640 media with was refreshed 48 hours prior to using cells for experiments with dHL-60 cells.

### Primary Polymorphonuclear Leukocyte (PMN) Isolation

Whole blood samples from a healthy donor screened by the American Red Cross were used for PMN isolation. The PMNs were separated using the density gradient reagent Polymorphprep (Progen) based on the manufacturer’s instructions. Briefly, 5 ml of Polymorphprep was added to a 15 ml conical tube, then 5 ml of human whole blood was added on top. The tubes containing the Polymorphprep and blood were centrifuged at room temperature at 500xg for 40 minutes to separate the blood components. The density gradients were carefully unloaded and the PMN fraction was collected. The PMNs were washed in sterile PBS and centrifuged to pellet at 400xg for 10 minutes. The pellet was resuspended in 2 ml of red blood cell (RBC) lysis buffer (eBioscience 1X RBC Lysis Buffer) for 30 seconds, then diluted in 10 ml serum free RPMI 1640 media to neutralize the RBC lysis buffer. PMNs were centrifuged at 300xg for 5 minutes and red blood cell lysis was repeated once more if there was visible red blood cell contamination of the pellet. PMNs were then resuspended in serum free RPMI media and counted using trypan blue to achieve a concentration of 10^6^ cells/mL.

### Microplate Assay to Evaluate Extracellular DNA (dHL-60 and PMN)

dHL-60 cells or primary PMNs were harvested and resuspended in serum free RPMI 1640 media at a concentration of 10^6^ cells/mL. 100 µL of the cell suspension was added to the wells of a black optical bottom 96 well plate (Thermo Scientific Nunc) to achieve a seeding density of 10^5^ cells/well. 20 µL of the designated nanoparticle treatment was added to each well, or PBS as a control in cells that received no nanoparticle treatment. In addition, blank wells with no cells containing 100 µL of RPMI media and 20 µL of the designated nanoparticle treatment were prepared in parallel to control for background Sytox fluorescence, especially from the PGM PLGA nanoparticle-associated extracellular DNA. In the prophylactic treatment experiments, cells were incubated for 30 minutes at 37°C and 5% CO_2_ prior to stimulation of NETosis with PMA or an equal volume of a vehicle in unstimulated groups. In the therapeutic treatment experiments, PMA or a vehicle was added immediately after plating the cells in each well and incubated for 0, 30, or 60 minutes before adding the designated nanoparticle treatment. The final concentration of PMA added to PMNs was 50 nM and to dHL-60s was 100 nM. Blank control wells with no cells received an equal volume of PMA or the vehicle. Cells were incubated for 4 hours at 37°C and 5% CO_2_. The extracellular DNA dye Sytox green (Invitrogen) was diluted in sterile PBS to 0.5 µM in the PMN experiments and 1 µM in dHL-60 experiments. 10% v/v of the diluted Sytox green dye was added to each well, to achieve a final concentration of 0.05 µM in PMN experiments and 0.1 µM in dHL-60 experiments. Cells incubated for 15 minutes, then the Sytox green fluorescence was read on a microplate reader at excitation and emission wavelengths of 504 nm and 523 nm, respectively.

### O-linked Glycosylation Fluorometric Assay

To quantify the relative mucin concentration in each nanoparticle formulation, the fluorescence of O-linked glycans was quantified using a previously developed assay.^54^ 50 µL aliquots of each nanoparticle formulation diluted 1:10 in ultrapure water were prepared. A standard curve of PGM was prepared using a stock solution of 5 mg/mL and serially diluted 1:2 in water 6 times. An aliquot of blank ultrapure water was also included as a standard. A solution with a final concentration of 0.6 M 2-cyanoacetamide (CNA) and 0.15 M sodium hydroxide (NaOH) was prepared. The CNA and NaOH solutions were mixed with 1 part CNA to 5 parts NaOH. A volume of 60µL of the CNA/NaOH solution was added to each 50 µL aliquot of the diluted nanoparticle sample or PGM standard and incubated on a heat block at 100°C for 30 minutes. 500 µL of a 0.6 M boric acid buffer (pH 8.0) was added to each sample or standard to quench the reaction. Samples and standards were added to the wells of a back 96 well plate and the fluorescence was quantified using a microplate reader with excitation and emission wavelengths of 336 and 383 nm, respectively. The concentration of PGM present in each formulation was calculated based on the fluorescence values of the standard curve. The final concentration of PGM that dHL-60 or PMN cells would be exposed was calculated based on the volumetric dilution of the nanoparticles used in subsequent *in vitro* experiments.

## Results and Discussion

Based on findings by ourselves and others that mucins strongly inhibit NET formation^12,16^, we sought to develop PLGA NP formulations that incorporate mucins to suppress NETosis while simultaneously retaining the ability to deliver therapeutics intracellularly. Following previous reports of amphiphilic proteins used as stabilizers for nanoemulsions^20^, we incorporated PGM into our emulsion via sonication to both stabilize the particles and associate to the particle surface (**Figure 1**). Typically, a stabilizer is added to prevent the emulsion droplets from coalescing before the organic solvent is removed, leaving behind solid PLGA particles that require washing to remove the surface associated stabilizers. As the commercially available PGM contains bacterial contaminants and insoluble agents, we adapted our NP synthesis protocol to include an additional centrifuge step after solvent evaporation to pellet any debris. Afterward, contaminants or large aggregates were removed via sterile filtration. This approach could be used to stabilize other oil in water nanoemulsions, including different polymer and lipid formulations^21,22^. To test the ability of our NPs to inhibit NETosis *in vitro*, we used microplate-based assays to quantify the extracellular DNA (**Figure 1**). As a measure of NETosis, the cells were stained with Sytox green, a membrane impermeable DNA dye, and the fluorescence was quantified using a plate reader. Similar plate reader assays have been employed across many studies to quantify NETosis using Sytox dyes^23–25^. Since the PGM itself contains residual DNA, a plate reader assay format is useful in allowing us to subtract out the fluorescence values from cell-free controls to ensure that we are excluding PGM-associated DNA and quantifying only changes in NET-associated DNA fluorescence.

**Figure 1.**
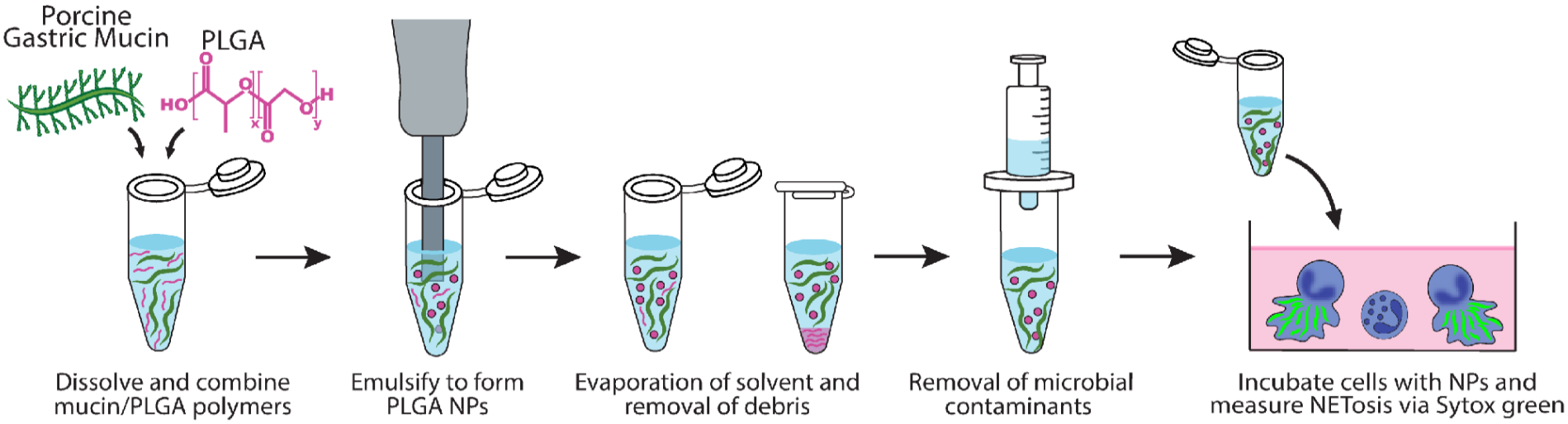
Schematic overview of the PGM-PLGA NP formulation and *in vitro* evaluation of NETosis in subsequent figures.

### PGM-PLGA Nanoparticle Characterization

We formulated particles with varying concentrations of PGM (0, 1, 2.5, or 5 mg/ml) in the aqueous phase both to optimize particle properties and to later find the most efficacious concentration for NETosis inhibition^26^. Based on the results of dynamic light scattering analysis, the size of the particles increased with the addition of PGM (**Figure 2A**). Additionally, the zeta potential of the particles became less negative upon the addition of the PGM (**Figure 2B**). As expected, the polydispersity index also increased with the addition of mucins (**Figure 2C**). The PGM-PLGA NP synthesis process may also cause some loss of mucins or entrapment in the particle core. To quantify mucin concentrations after particle formation, we used a fluorometric assay to determine the relative concentration of O-linked glycans present in each of the 0, 1, 2.5 and 5 mg/ml PGM-PLGA NP formulations. We found PGM concentration in each formulation increased proportionally with the initial mucin input concentration (**Figure 2D**). These NPs were then further studied for their ability to suppress NETosis and deliver model drug cargoes

**Figure 2.**
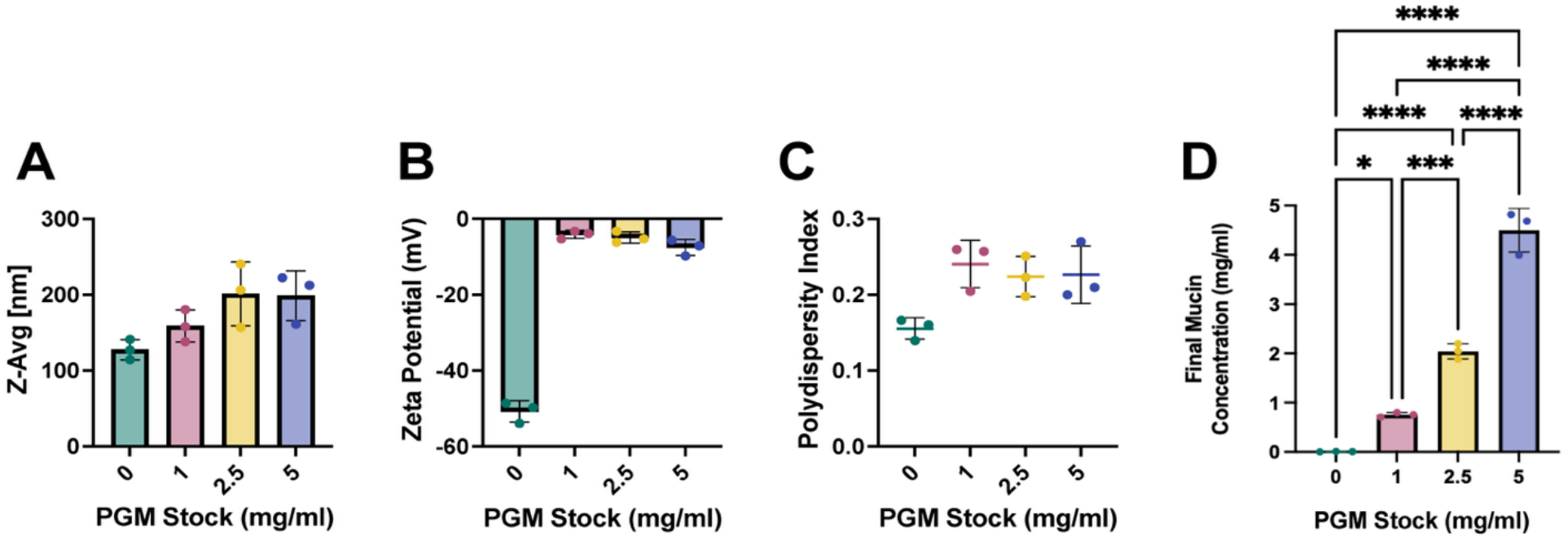
Characterization of PGM-PLGA NPs. PGM-PLGA NPs were formulated with increasing concentrations of PGM with stock solution concentrations of 0 (PLGA-only NPs), 1, 2.5, or 5 mg/ml PGM. **A)** Diameter, **B)** zeta potential and **C)** polydispersity index of each NP formulation using dynamic light scattering (DLS). **D)** The relative mucin concentration of each NP formulation was determined using a fluorometric assay to detect O-linked glycans.

### PGM-PLGA NPs Inhibit NETosis in dHL-60 Cells

After characterizing the particle’s physiochemical properties, we sought to test our PGM-PLGA particles’ ability to inhibit NETosis. The mechanism underlying PGM NETosis inhibition is thought to be dependent on sialic acid terminated glycans. Sialylated mucins can bind to inhibitory cell surface receptors called sialic acid-binding immunoglobulin-like lectins (Siglecs), leading to downregulation of ROS production in the neutrophils^16^. Therefore, for our particles to inhibit NETosis, we anticipated surface exposure of the mucins will be required to be engaged by neutrophils via Siglec binding. Based on well understood biophysical principles of protein adsorption to exposed nanoparticle surfaces^27,28^, the particles with surface associated mucins could have an interface with a high local concentration of mucins as compared to their concentration in the bulk solution. However, the confirmation of surface-associated mucins at the particle interface could also change accessibility of mucin-associated glycans for Siglec binding by neutrophils. Thus, we reasoned NETosis inhibition mediated by mucins associated to a nanoparticle surface may differ from that of an aqueous mucin solution at a similar concentration.

We assessed the ability of PGM-PLGA NPs to inhibit NETosis at various PGM concentrations in our plate reader assay with differentiated HL-60 cells. HL-60 cells are a promyelocytic cell line that differentiate into a neutrophil-like state upon exposure to low levels of dimethyl sulfoxide (DMSO) for 5-7 days. These differentiated HL-60 (dHL-60) cells are a widely used neutrophil-like model cell line. dHL-60 cells mimic many neutrophil phenotypes, including the ability to undergo NETosis. We administered the NPs 30 minutes before stimulating the dHL-60 cells with PMA to induce NETosis (**Figure 3A**). PMA is a well-characterized inducer of NETosis and does not cause apoptosis^29^. After the addition of PMA, we see a sharp increase in the Sytox fluorescence intensity of ∼2.5 fold that is indicative of successful induction of NETosis, as shown where unstimulated dHL-60 cells have a low RFU of ∼5000 (**Figure 3C**) while PMA stimulated jump to ∼12500 RFU (**Figure 3B**). When PLGA particles without PGM (0 mg/ml) are added, we see an increase in RFU up to ∼20000 in the PMA stimulated cells, significantly higher than the PMA stimulated cells with no NP treatment (**Figure 3B**). This suggests that PLGA itself is a potent inducer of NETosis in activated neutrophils, matching results found by Brannon et al^7^. On the contrary, when dHL-60 were dosed with the 1, 2.5, and 5 mg/ml PGM-PLGA NP groups, we see a reduction in RFU, with 2.5 and 5 mg/ml PGM-PLGA NP treatment returning the total NETosis back to the ∼5000 RFU baseline of unstimulated cells (**Figure 3B**). Even with unstimulated cells, where we see no significant difference in the RFU of the NP, 0 mg/ml, 1 mg/ml groups, the 2.5 and 5 mg/ml PGM-PLGA NP groups reduce the RFU down to an almost undetectable intensity, meaning viability is preserved and spontaneous NETosis is reduced (**Figure 3C**).

**Figure 3.**
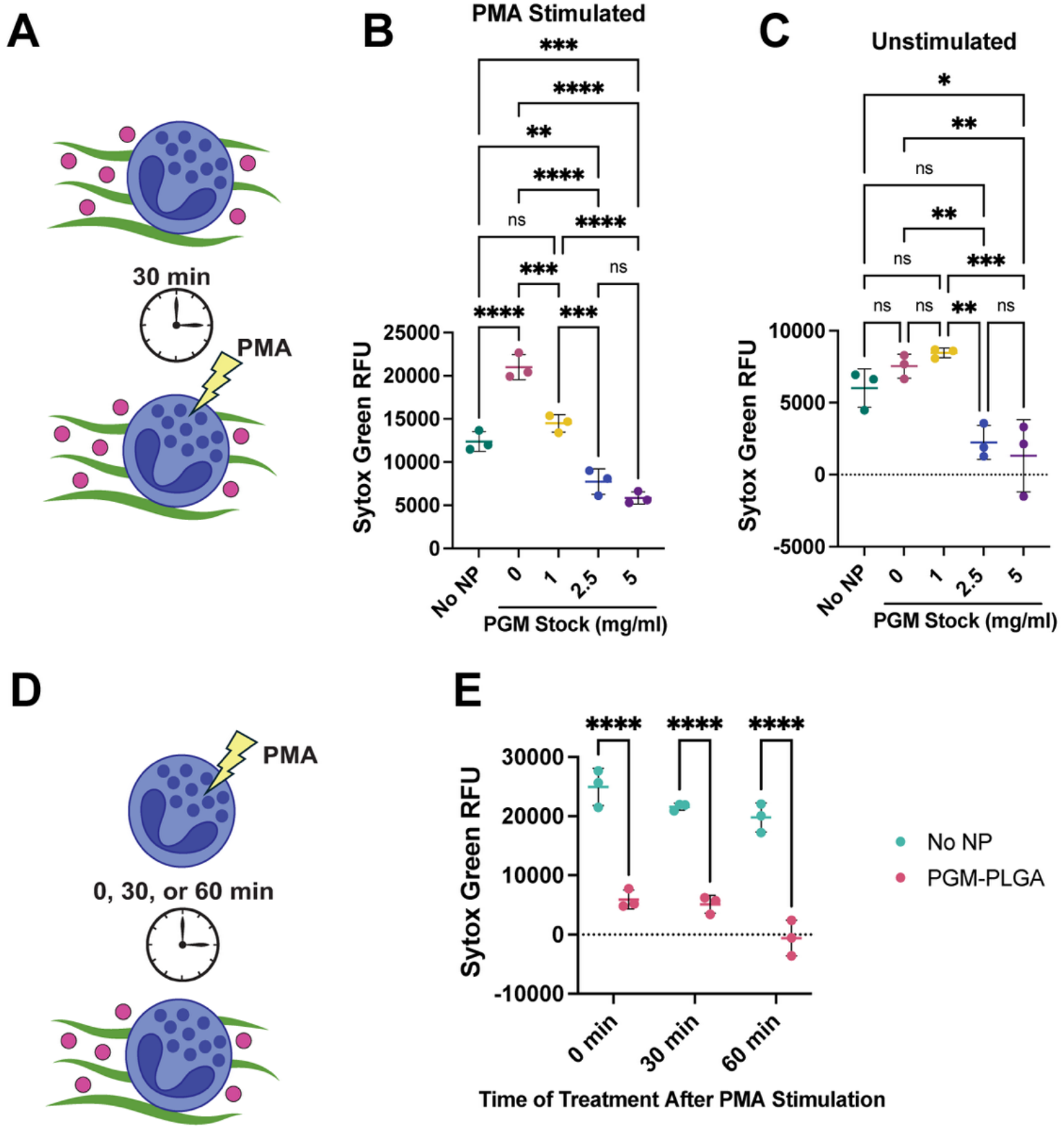
PGM-PLGA NPs Ameliorate NETosis in dHL-60 Cells. **A)** Schematic of PGM-PLGA NP application in experiments to test prophylactic efficacy in preventing NETosis. The Sytox green fluorescence of dHL-60 cells treated with NPs formulated with a PGM stock solution of 0 (PLGA-only NPs), 1, 2.5, or 5 mg/ml compared to control wells treated with a vehicle (no NP) was measured using a microplate-based assay. **B)** All dHL-60 cells were stimulated with 100 nM PMA to induce NETosis. **C)** All dHL-60 cells were left unstimulated and treated with PBS instead of PMA to evaluate spontaneous NETosis. **E)** Schematic of PGM-PLGA NP application in experiments to test therapeutic efficacy in stopping NETosis after PMA stimulation. **F)** Sytox green fluorescence of dHL-60 cells treated with NPs formulated with a PGM stock solution of 5 mg/ml at 0 (co-treatment), 30, or 60 minutes following PMA stimulation compared to control wells treated with a vehicle (no NP) using a microplate-based assay. * p < 0.05, ** p < 0.01, *** p < 0.001, **** p < 0.0001, ns = not significant. Statistical tests performed in B-D) are one-way ANOVAs with Tukey’s multiple comparisons test. In F), the statistical test performed was a two-way ANOVA with Šidák’s multiple comparisons test.

We decided to pursue the 5 mg/ml PGM-PLGA NP formulation as the strongest NETosis inhibitor and compare it to the 0 mg/ml NP group. Henceforth, PLGA NPs refer to the 0 mg/ml group and PGM-PLGA NP refer to the 5 mg/ml PGM-PLGA NP group. To give more insight into our PGM-PLGA NP formulation, we evaluated if PGM was strongly associated with PLGA NPs by washing the NPs multiple times before treating dHL-60 cells. We found that PGM-PLGA NPs subjected to washing lost the ability to inhibit NETosis, showing the same effect as the PLGA NPs in increasing Sytox green fluorescence compared to the no NP control (**Figure S1**).

Our next *in vitro* experimental design took into consideration that NP therapeutics for inflammatory conditions may interact with neutrophils already primed to undergo NETosis. NETosis takes ∼1-4 hours for the cell to fully lyse once the pathway is induced ^30^. Therefore, we modified our experimental scheme to stimulate the dHL-60 cells with PMA before adding PGM-PLGA NPs, priming them to undergo NETosis before any potential inhibition could occur. We dosed PGM-PLGA NPs both 30 minutes and 1 hour after PMA stimulation (**Figure 3D, E**). To ensure NETosis could occur, we continued the PMA stimulation for a total of four hours^30^. PGM-PLGA NPs prevented primed dHL-60 cells from releasing NETs, as evidenced by the significant decrease in Sytox green RFU from ∼20000 in the PLGA NP treated groups to less than 5000 RFU (**Figure 3E**).

### PGM-PLGA NPs Inhibit NETosis in Primary Human Polymorphonuclear Leukocytes

Though dHL-60 cells are an excellent model of neutrophils, primary human polymorphonuclear leukocytes (PMNs) are the most physiologically relevant *in vitro* model. Therefore, we tested the ability of our PGM-PLGA NPs to inhibit NETosis in primary PMNs isolated from the blood of healthy human donors. Our PMA stimulation and NP intervention scheme for PMN’s followed that of our dHL-60 cells. First, we treated PMNs with PLGA or PGM-PLGA NPs for 30 minutes prior to stimulation to induce NETosis with PMA or a vehicle control in the unstimulated groups. PLGA NPs again induced greater NETosis compared to the no NP control in both the PMA stimulated and unstimulated PMNs, significantly increasing the Sytox green RFU (**Figure 4A, C**). In comparison, the PGM-PLGA NPs significantly decreased the Sytox green RFU in both the PMA stimulated and unstimulated groups, indicating a large reduction in NETosis (**Figure 4A, C**). This effect can be visualized with brightfield images seen in **Figure 4B**, where PMA stimulated cells with PLGA nanoparticles and PGM-PLGA nanoparticles have distinct morphologies. In the no NP and PLGA NP images, PMNs undergoing NETosis appear large, flattened, and more transparent as the cell membrane lyses and the chromatin NET complex is released. In the PGM-PLGA NP treated PMNs, the cells appear smaller in size, more rounded, and refractile due to the cell membranes being intact. Overall, we see that PGM-PLGA NPs can successfully reduce PLGA NP exacerbated NETosis in both stimulated and unstimulated primary human PMNs. We also repeated our experiment in which PMNs were primed with PMA for 1 hour prior to treatment with the PGM-PLGA NPs (**Figure 4D**). PGM-PLGA particles were able to similarly inhibit NETosis of PMNs, suggesting that the PGM-PLGA NPs may successfully reduce NETosis in inflammatory *in vivo* environments.

**Figure 4.**
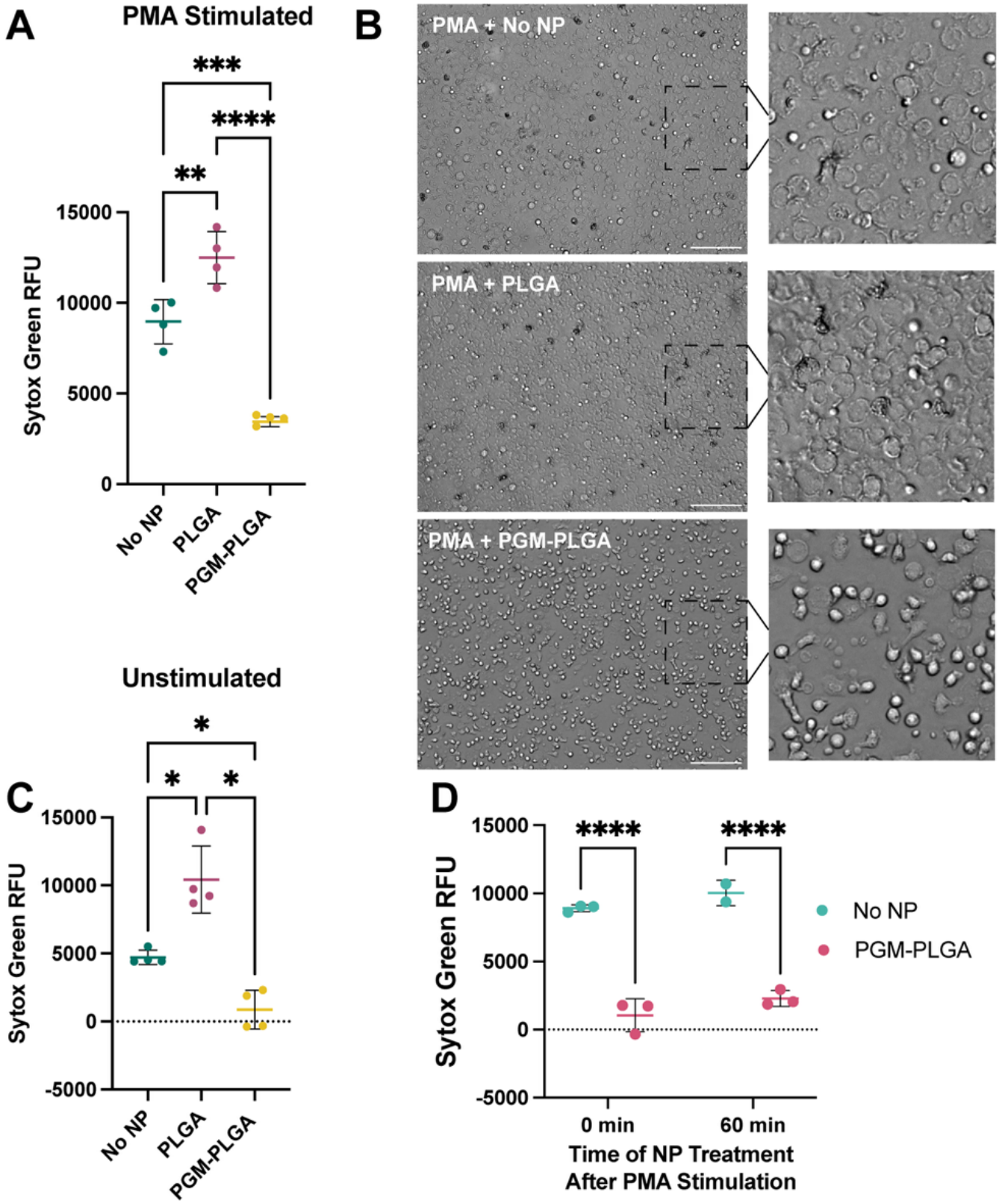
PGM-PLGA NPs Reduce NETosis in Primary Human PMNs. **A)** The Sytox green fluorescence of dHL-60 cells treated with NPs formulated with a PGM stock solution of 0 (PLGA-only NPs) or 5 mg/ml (PGM-PLGA) compared to control wells treated with a vehicle (no NP) was measured using a microplate-based assay. All PMNs were stimulated with 100 nM PMA to induce NETosis. **B)** Brightfield images (20x magnification) of PMA-stimulated PMNs following 4 hours of treatment with a vehicle (no NP), PLGA NPs, or PGM-PLGA NPs. Scale bars, 100 µm. Zoomed in portions to better show cell morphology are also shown beside each image. **C)** dHL-60 cells were left unstimulated and treated with PBS instead of PMA to evaluate spontaneous NETosis using the same Sytox-green based microplate assay and NP treatments. **D)** Sytox green fluorescence of dHL-60 cells treated with NPs formulated with a PGM stock solution of 5 mg/ml at 0 (co-treatment) or 60 minutes following PMA stimulation compared to control wells treated with a vehicle (no NP) using a microplate-based assay. * p < 0.05, ** p < 0.01, *** p < 0.001, **** p < 0.0001. Statistical tests performed in A) and C) are one-way ANOVAs with Tukey’s multiple comparisons test. In D), the statistical test performed was a two-way ANOVA with Šidák’s multiple comparisons test.

### PGM-PLGA NPs Have Promising Drug Delivery Characteristics

PLGA NPs are a well-characterized modality for drug delivery, with the ability to encapsulate and deliver both protein and small molecule therapeutics^20,26^. While PLGA NPs could serve as an extracellular depot of therapeutics, they may also be internalized for intracellular delivery^31,32^. However, it is unknown whether interactions of cells with PGM to inhibit NETosis would reduce intracellular delivery of PLGA NPs. Previous studies indicate that mucins may reduce phagocytosis of bacteria by PMNs^33,34^. Furthermore, while reducing NETosis can offer substantial therapeutic benefit, NET inhibition alone hasn’t proven to be an all-encompassing treatment for inflammatory conditions^35^. Therefore, retention of the intracellular drug delivery capabilities of PGM-PLGA NPs is crucial to consider for maximizing future therapeutic uses for neutrophilic inflammatory diseases.

We performed several well-established drug delivery assays to confirm NP stability, drug encapsulation, and cellular uptake. We performed NTA to obtain a measure of the nanoparticle concentration of our PLGA and PGM-PLGA groups. As expected, using PGM as a stabilizer increased the number of particles in solution from an average of 4.5 × 10^11^ particles per ml for PLGA to 1.0 × 10^12^ particles per ml for PGM-PLGA, suggesting there was less loss due to aggregation during the initial sonication, drying, and filtering phases (**Figure 5A**). We used the fluorescent dye DiD, a far-red fluorescing hydrophobic molecule, to act as a reporter system for molecule encapsulation and cellular uptake. We loaded both PLGA and PGM-PLGA NPs with DiD by dissolving it alongside PLGA in the chloroform phase of our nanoemulsion. Based on encapsulation analysis, PGM-PLGA particles also have significantly increased DiD encapsulation up to 1 µg/ml (16% encapsulation efficiency), likely due to the increase in particle count and improved stabilization (**Figure 5B**). When comparing the concentration of DiD against the concentration of particles, no significant difference was observed between PLGA and PGM-PLGA, suggesting the amount of DiD per particle didn’t change with the addition of PGM. To evaluate stability, we measured particle size over time in PBS using DLS, with both PLGA and PGM-PLGA showing no change in size over 12 hours in PBS (**Figure 5C**). Likely, PLGA particles are stabilized by their highly negative charge inducing electrostatic repulsion^36^.

**Figure 5.**
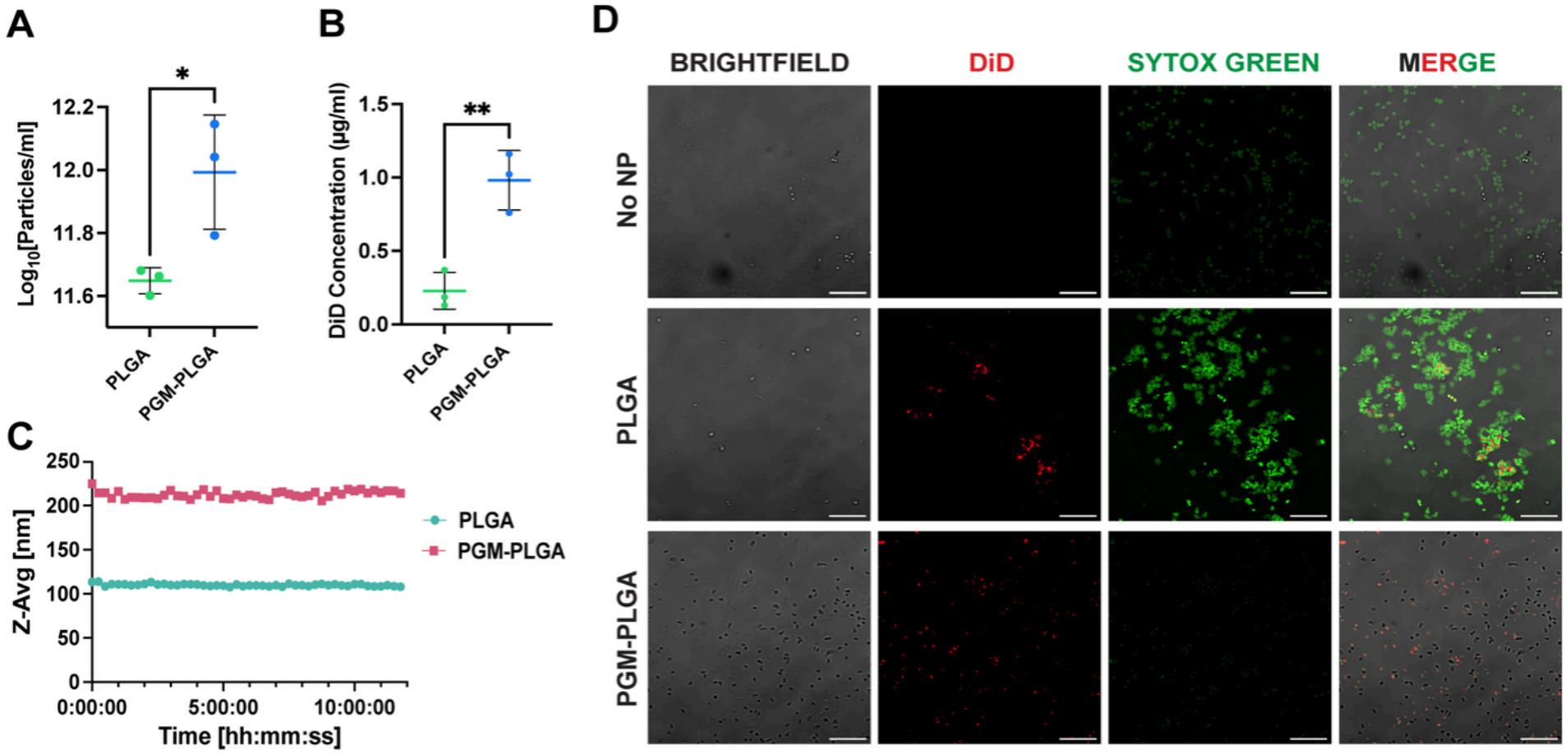
PGM-PLGA NPs Successfully Deliver Cargo to dHL-60 Cells. **A)** NP concentration in PLGA and PGM-PLGA NP formulations was evaluated using DLS. **B)** The encapsulation efficiency of each NP formulation was assessed by measuring the internal DiD concentration. **C)** The stability of PLGA and PGM-PLGA NPs in PBS over the course of 12 hours. **D)** Confocal microscopy images (10x magnification) of PMA-stimulated dHL-60 cells treated with a vehicle (no NP), PLGA NPs, or NPs formulated with a PGM stock of 5 mg/ml (PGM-PLGA). All NPs were loaded with DiD to evaluate uptake after 4 hours of incubation. In addition, Sytox green staining was used to visualize NETosis. Scale bars, 50 µm. * p < 0.05, ** p < 0.01. Statistical tests performed in A) and B) are unpaired t-tests.

After confirming encapsulation, PMA stimulated dHL-60 cells were imaged to assess NETosis nanoparticle uptake (**Figure 5D**). There are bright, large areas of Sytox green visible in both the no NP and PLGA NP treated groups stimulated with PMA, indicating cells were undergoing NETosis. Interestingly, in the PLGA NP treated cells, there is localization of the DiD loaded PLGA particles to the NETs. When comparing the PGM-PLGA group to these groups, we can see that there is a large reduction in intensity and smaller areas of Sytox green that suggest reduced NETosis. There is still some faint staining of Sytox green in the PGM-PLGA images, which we attribute to incomplete washing away of trace amounts of Sytox dye after staining the live cells prior to fixation (live cells were washed three times before 4% PFA fixation). The PGM-PLGA NP treated dHL-60 cells appear to be viable, similar to the PMN images in **Figure 4B**. The cells that have adhered to the bottom of the glass slide are in focus in **Figure 5D**, but in the brightfield images we see many other intact, viable cells in nearby z-planes in the PGM-PLGA NP treatment group. In comparison, in both the no NP and PLGA NP brightfield images we see very few cells in other z-planes, likely meaning most cells have undergone NETosis and adhered to the bottom of the glass slide. Moreover, DiD-loaded NPs were distributed across all cells and appear to be intracellular. This is a stark comparison to the PLGA NP treated group, in which DiD loaded NPs aggregate in large clusters associated with extracellular NETs. Ultimately, this may present a significant advantage of the PGM-PLGA NP formulation in enhancing intracellular drug delivery by inhibition of NETosis.

To confirm that the use of PGM in the PGM-PLGA NP formulation does not compromise cellular uptake, we performed flow cytometry of HL-60 cells dosed with DiD loaded particles. The viable cells in both the PLGA and PGM-PLGA NP treated groups exhibited 100% uptake at the same dosing used in all assays (**Figure S2**). In addition, there were no differences in the geometric mean DiD intensities between the live cells in the PLGA and PGM-PLGA NP treated groups in either the PMA stimulated or unstimulated groups. However, PMA stimulation significantly decreased geometric mean of the DiD signal, suggested activated cells have lowered uptake of these particles (**Figure S2**). Taken together, our results show that PGM stabilization does not compromise the intracellular drug delivery potential of PLGA NPs, and PGM-PLGA NPs provide improved particle retention and encapsulation efficiency.

Our results point towards an increase in functionality with mucin stabilization of PLGA NPs, including general particle stabilization and NETosis inhibition. Current research suggests that PLGA NPs exacerbate NETosis, corroborated by our findings^7^. PLGA induced NETosis could stem from nominal particulate properties that can destabilize membranes^37^, such as particle size and hydrophobicity, or from PLGA degradation into lactic acid, which can potentially induce NETosis^38^. However, these interactions are not well understood and further research into how PLGA induces NETosis is warranted. Herein, we built upon previous findings from our group and others that mucins can suppress NETosis to develop PGM-PLGA NPs. We found that PGM-PLGA NPs successfully reduce NETosis and retain intracellular drug delivery capabilities. The use case for the PGM-PLGA NPs could fit into a dual inflammation targeting regime, where particles deliver anti-inflammatory molecules while inhibiting NETosis, or by using neutrophils to backpack to sites of inflammation^39^. Future studies to further characterize the PGM-PLGA NPs could include dosing response to investigate cellular uptake further and active drug loading to investigate further reduction in neutrophilic inflammation^40^. Beyond particles, the other FDA-approved modalities of PLGA-based drug delivery include in situ forming gels and solid implants. Our approach could be adapted for PLGA-based solid implants by grafting mucins to the material, as prior work has developed processes to coat several different material surfaces with mucins^15^. Ultimately, bioactive and inspired materials like these may help push immunomodulatory treatments forward.

## Acknowledgements

This project was funded by the Cystic Fibrosis Foundation (DUNCAN24G0) and National Institutes of Health (R01 HL160540, F31HL176146).

## Author contributions

A.B., O.T., and G.A.D. conceptualized the project and designed experiments. A.B., O.T., and A. J. conducted experiments and led data analysis. A.B., O.T., and G.A.D. wrote the article. All authors have given approval to the final version of the manuscript.

## Data Availability

Data can be made available upon request.

