## Supplementary Information for "Mucin-laden PLGA nanoparticles inhibit neutrophil extracellular trap formation"

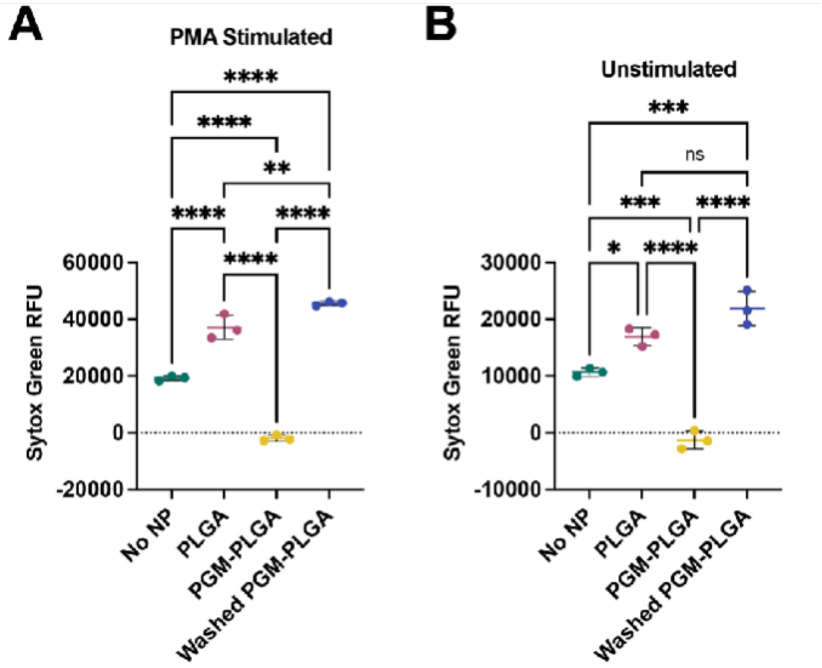


**Figure S1.** In order to determine if mucins were bound to PGM-PLGA NPs, the PGM-PLGA NPs were washed three times in sterile PBS and resuspended to approximately the same volume afterwards. The washed PGM-PLGA NPs were applied to dHL-60 cells and the Sytox green fluorescence was quantified using a microplate-based assay. dHL-60 cells treated with the unwashed PGM-PGLA NPs, PLGA-only NPs, and no NPs. **A)** dHL-60 cells stimulated with 100 nM PMA to induce NETosis. **B)** dHL-60 cells unstimulated with PBS. * $p < 0.05$, ** $p < 0.01$, *** $p < 0.001$, **** $p < 0.0001$, ns = not significant. Statistical tests performed in A) and B) are one-way ANOVAs with Tukey's multiple comparisons test.

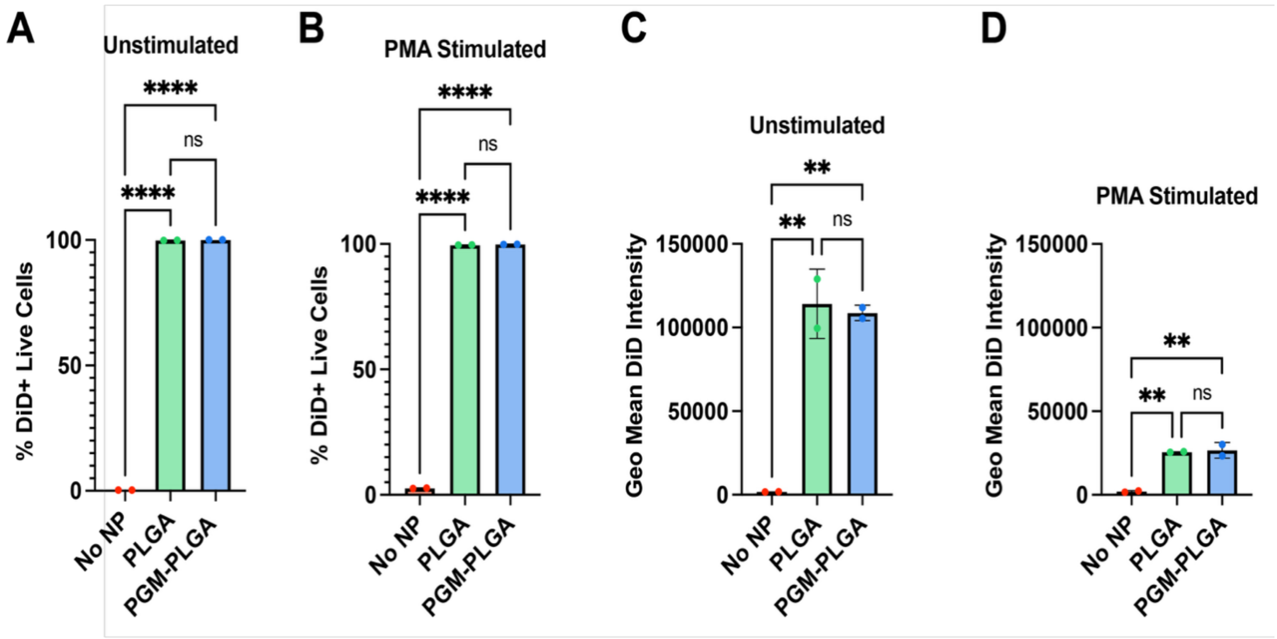


**Figure S2.** Flow cytometry of dHL-60 cells treated with the same NP formulations loaded with DiD was performed to evaluate uptake efficacy. Cells were incubated with NP or vehicle treatments for 30 minutes prior to adding PBS (unstimulated) or 100 nM PMA (stimulated) and incubation for an additional 1 hour. **A)** Results showing the percentage of live DiD+ cells in the unstimulated and **B)** PMA-stimulated cells. The geometric mean DiD intensity of the live DiD+ live cell population is reported in the **C)** unstimulated and **D)** PMA stimulated cells. * $p < 0.05$, ** $p < 0.01$, **** $p < 0.0001$, ns = not significant. Statistical tests performed in A-D) are one-way ANOVAs with Tukey's multiple comparisons test.
